# Polymer skulls with integrated transparent electrode arrays for cortex-wide opto-electrophysiological recordings

**DOI:** 10.1101/2021.11.13.468490

**Authors:** Preston D. Donaldson, Zahra S. Navabi, Russell E. Carter, Skylar M. L. Fausner, Leila Ghanbari, Timothy J. Ebner, Sarah L. Swisher, Suhasa B. Kodandaramaiah

**Affiliations:** Department of Electrical and Computer Engineering, University of Minnesota Twin Cities; Department of Mechanical Engineering, University of Minnesota Twin Cities; Department of Neuroscience, University of Minnesota, Twin Cities; Department of Biomedical Engineering, University of Minnesota Twin Cities

## Abstract

Electrophysiological and optical imaging provide complementary neural sensing capabilities – electrophysiological recordings have the highest temporal resolution, while optical imaging allows recording the activities of genetically defined populations at high spatial resolution. Combining these complementary, yet orthogonal modalities to perform simultaneous large-scale, multimodal sensing of neural activity across multiple brain regions would be very powerful. Here we show that transparent, inkjet-printed electrocorticography (ECoG) electrode arrays can be seamlessly integrated with morphologically conformant transparent polymer skulls for multimodal recordings across the cortex. These ‘eSee-Shells’ were implanted on transgenic mice expressing the Ca^2+^ indicator GCaMP6f in cortical excitatory cells and provided a robust opto-electrophysiological interface for over 100 days. eSee-Shells enable simultaneous mesoscale Ca^2+^ imaging and ECoG acquisition under anesthesia as well as in awake animals presented with sensory stimuli. eSee-Shells further show sufficient clarity and transparency to observe single-cell Ca^2+^ signals directly below the electrodes and interconnects. Simultaneous multimodal measurement of cortical dynamics reveals changes in both ECoG and Ca^2+^ signals that depend on the behavioral state.

## INTRODUCTION

Many sensorimotor and cognitive behaviors require the interaction of activities in widespread, disparate brain regions. Neuroscientists have traditionally focused on understanding the roles that single brain regions play in mediating behavior. However, much less is known about how neuronal activities across regions are coordinated, and how that coordination dynamically changes as a function of behavioral state. A wealth of recent work investigating cortical dynamics across large cortical regions has revealed complex dynamics associated with a variety of behaviors^1–3^. We know that ongoing activity is attuned with locomotion^4,5^, and these dynamics are altered as the cognitive complexity of the task changes^6^. There are also specific, distributed activities during goal-directed behaviors^7^ and decision making^8^.

Such studies have typically used transparent windows for calcium (Ca^2+^) imaging which allow high-resolution cellular mapping of activities from microcircuits^9^. The advent of large cranial windows^10–12^ allows either mesoscale mapping of Ca^2+^ activity or random access measurement of cellular activity from smaller fields of view (FOV). As large-scale ongoing activity modulates single cells within microcircuits^13^, it is advantageous to simultaneously monitor both cellular and mesoscale activity. To do so, approaches for simultaneous mesoscale imaging and 2-photon (2P) cellular resolution imaging^14^ or measuring cellular-like activities from across the cortex^15^ have been developed.

However, Ca^2+^ imaging is limited in temporal resolution and unable to capture the full range of frequency-specific information essential for many sensorimotor and cognitive behaviors^16–18^. Combining multi-scale Ca^2+^ imaging with high temporal resolution electrophysiological recordings would be very powerful. Recent advancements in transparent electrocorticography (ECoG) electrode arrays^19–29^ allow simultaneous Ca^2+^ imaging and ECoG acquisition. To date, these efforts have largely focused on small FOVs, encompassing single brain regions. Extending multimodal monitoring to cortex-wide neural activity has yet to be realized.

In this work, we demonstrate the seamless integration of inkjet-printed, transparent electrode arrays onto transparent polymer skulls^11^, for simultaneous, multimodal neural sensing over a large fraction of the mouse dorsal cerebral cortex. The functionalized polymer skulls, or ‘eSee-Shells’, consist of 10 electrodes spread over a window with a field of view of ~45 mm^2^. We report long-term multimodal recordings lasting months, with the ability to perform mesoscale or cellular-resolution Ca^2+^ imaging simultaneously with ECoG acquisition.

## RESULTS

### eSee-Shell design

A device that enables simultaneous electrophysiology and optical imaging of neural activity across much of the mouse dorsal cortex requires several key functionalities. First, the device needs to conform to the complex 3D surface of the brain. Second, the device needs to be transparent and have sufficient optical clarity to allow high spatial resolution optical imaging. Third, this transparent interface needs to integrate electrodes that are both transparent and flexible to allow conformity to the brain surface. We recently developed transparent polymer skulls that allow sub-cellular resolution optical imaging across most of the dorsal cortex of the mouse^11^. Further, we demonstrated that the polyethylene terephthalate (PET) film, the transparent window material in the polymer skulls, can be used as a substrate to pattern inkjet-printed transparent electrodes and interconnects^30^. Combining these capabilities, we developed the eSee-Shell, a fully-integrated transparent polymer skull with 10 inkjet-printed ECoG electrodes (**Fig. 1**).

**Figure 1:**
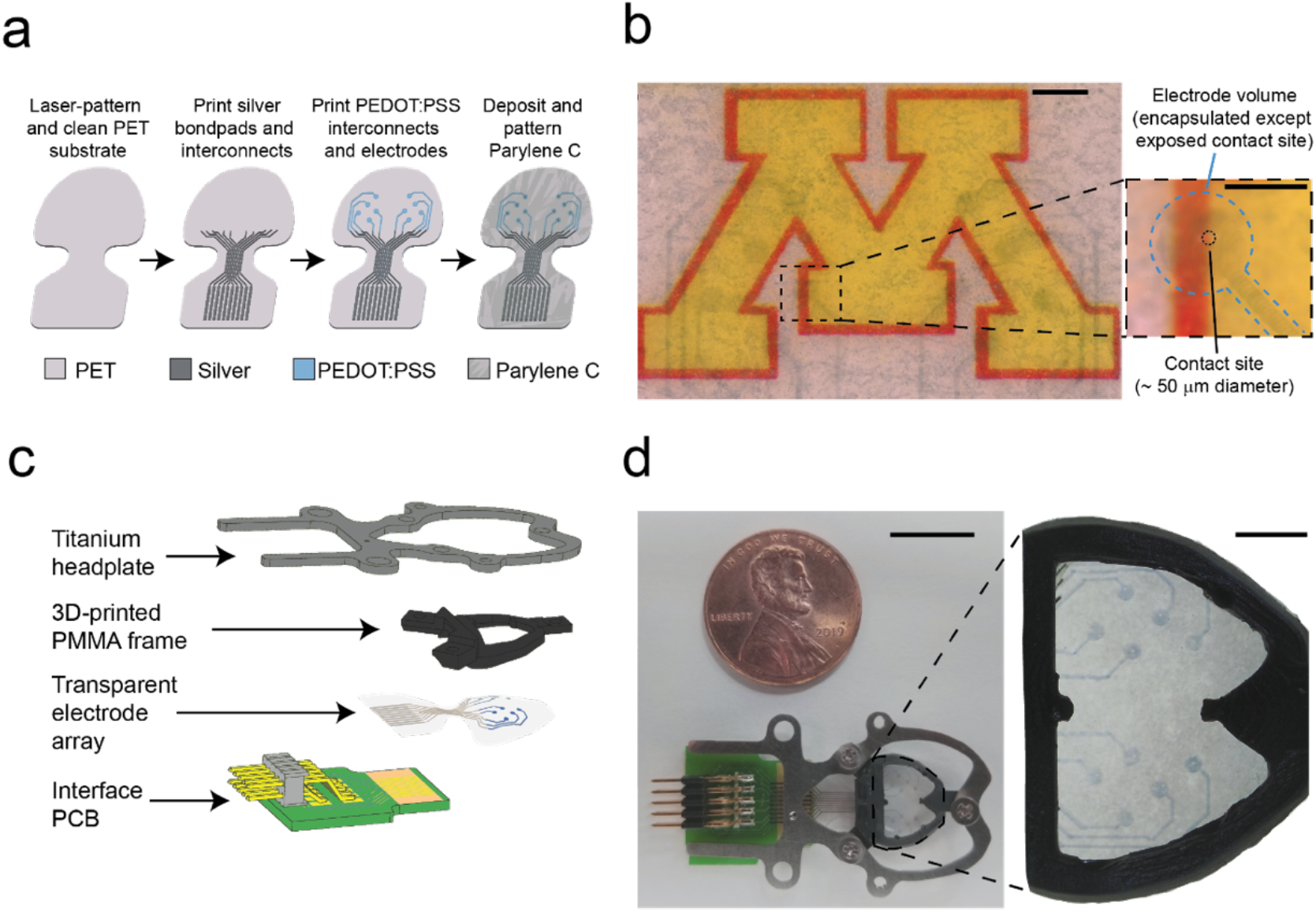
eSee-Shells. **(a)** Schematic diagram of electrode array fabrication process on planar PET substrate. **(b)** Left: Magnified photomicrograph of transparent electrode array on PET substrate overlaid on the University of Minnesota logo. Scale bar indicates 1 mm. Right: Zoomed-in photomicrograph of a single electrode, with a dashed line to indicate the border of the transparent electrode and the smaller circle to indicates the contact site. Scale bar indicates 500 μm. **(c)** Schematic of eSee-Shell components. **(d)** Left: Photograph of fully assembled eSee-Shell. Scale bar indicates 1 cm. Right: Zoomed-in photograph of the transparent electrode array bonded into a frame. Scale bar indicates 2 mm.

The electrodes and interconnects within the window area where transparency is required are composed of poly(3,4-ethylenedioxythiophene) polystyrene sulfonate (PEDOT:PSS), a transparent biocompatible conductor that is often used to reduce the interfacial impedance of neural interfaces^31,32^. Near the edge of the window area, the PEDOT:PSS interconnects contact inkjet-printed silver interconnects, which route the signals to bond pads for interfacing with a custom printed circuit board (PCB). The silver interconnects reduce the overall channel impedance as they are more conductive than PEDOT:PSS, and they are placed outside of the window area so as not to affect transparency. The entire array is then encapsulated with a biocompatible insulator, Parylene-C, and contact openings are etched over the electrodes. Because the silver interconnects are fully encapsulated and do not come into direct contact with the brain, there is little risk of neurotoxic responses to silver during chronic implantation.

The overall process for patterning and insulating the electrodes is illustrated in **Figure 1a**. Briefly, PET substrates were laser patterned to produce the outline of the electrode array and alignment features for later bonding to the window frame. Silver interconnects and bond pads were then patterned by inkjet printing, followed by inkjet printing of the PEDOT:PSS electrodes (diameter ~500 μm) and interconnects. The devices were encapsulated in Parylene-C, and then an oxygen plasma etch was used to create electrode contact openings in the Parylene-C. The unique electrode geometry used in eSee-Shells creates a low-impedance and transparent interface by using a large volume of encapsulated PEDOT:PSS spread laterally around a small ~50 μm contact site. The extra volume of PEDOT:PSS surrounding the contact site acts to reduce the channel impedance^30^, without diminishing transparency with the use of thicker PEDOT:PSS films. A photomicrograph of a completed electrode array is shown in **Figure 1b**. After completion of the electrode patterning, the functionalized PET film was bonded to the interface PCB using anisotropic conductive film (ACF). The functionalized PET film was then epoxy bonded to the polymethyl methacrylate (PMMA) frame of the polymer skull, as described previously^12,33^, to complete the eSee-Shell (**Fig. 1c-d**).

### Benchtop performance of the eSee-Shell electrodes

The electrode-electrolyte impedance of the eSee-Shell arrays in saline was characterized using electrochemical impedance spectroscopy (EIS). The average interfacial impedance spectrum for the ten electrodes of a single eSee-Shell array prior to bonding into the PMMA frame is shown in **Figure 2a**. The impedance spectrum reveals a low impedance at high frequencies (>1 kHz); this is typical of electrode interfaces and mostly due to the series resistance of the PEDOT:PSS interconnects. Higher impedances at low frequencies are typical of electrode-electrolyte interfaces^34^, and the impedance approaches 200 kΩ at 1 Hz for our PEDOT:PSS microelectrodes. This is much lower than previously reported transparent electrodes made with graphene^20,35^ or indium tin oxide (ITO)^27,29^, while also allowing smaller contact sites to be used. This large reduction in impedance at low frequencies results from the large volume of encapsulated PEDOT:PSS that can participate in signal transduction because of the volumetric capacitance of PEDOT:PSS films^30,36,37^. The eSee-Shell electrode arrays are robust and flexible enough to be deformed and bonded into the PMMA frame without affecting channel impedances. After bonding 6 microelectrode arrays into PMMA frames, only 1 out of 60 channels failed, and the average impedance of the remaining functional channels did not differ from before bonding (**Fig. 2b**).

**Figure 2:**
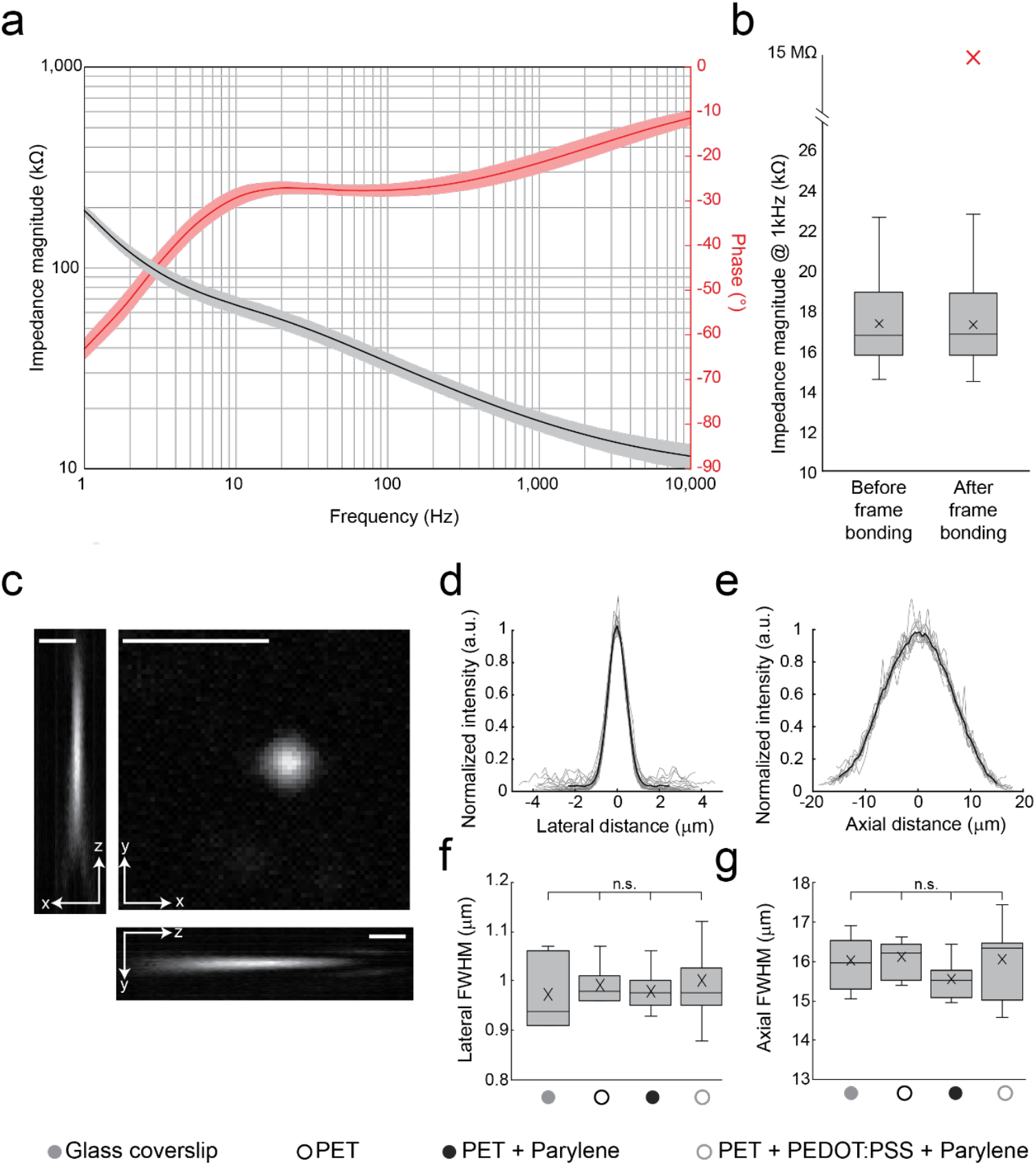
Benchtop characterizations of eSee-Shells. **(a)** Impedance spectra of printed PEDOT:PSS electrodes prior to bonding onto window frames (n = 60 electrodes). Black and red lines show impedance magnitude and phase, respectively, and shaded regions indicate standard deviation. **(b)** Impedance magnitude of electrodes at 1 kHz before and after being bonded onto window frames. Red ‘X’ indicates outlier (n = 60 electrodes). **(c)** Orthogonal sections of the point spread function (PSF) of a 0.1 μm fluorescent bead imaged with a confocal microscope through an electrode. Scale bars indicate 5 μm. **(d,e)** Lateral (n = 22) and axial (n = 12) intensity profiles of fluorescent beads imaged through electrodes. Grey lines indicate individual bead profiles, black lines indicate average profile. **(f,g)** Distributions of lateral and axial PSF full-widths at half maximum (FWHMs) when imaged through various material stacks. In all box-and-whisker plots, ‘x’ indicates mean, horizontal line indicates median, box edges indicate upper and lower quartiles, and whiskers indicate maximum and minimum values.

Another important characteristic of transparent electrodes is the resolution that can be attained when imaging through them. PET alone has a similar optical resolution to glass coverslips^11^. In eSee-Shells, the PET is the substrate for the PEDOT:PSS electrodes and interconnects, followed by an encapsulating layer of Parylene-C. Stacks of PET, PEDOT:PSS, and Parylene-C similar to those used in eSee-Shells transmit 80% and 77% of blue and green light, respectively, with ~2% or less haziness throughout the visible spectrum. To evaluate the resolution of imaging through the eSee-Shells, point spread functions (PSFs) of 0.1 μm yellow-green fluorescent beads dispersed in agar were measured with 1-photon (1P) confocal microscopy through the various film stacks. A representative image of a single PSF bead is shown in **Figure 2c**. For each bead, we measured the intensity profile along the lateral and axial dimensions and the intensities exhibited a Gaussian profile for both directions (**Fig. 2d,e**). We observed isotropic PSFs in the lateral plane, but as expected, the axial PSF is much wider than the lateral PSF because of additional contributions from out-of-focus light above and below the focal plane (which can be reduced in practice by using multiphoton microscopy or other advanced imaging methods). The full-width at half maximum (FWHM) values obtained from the Gaussian curves fit to the average normalized intensity reveal no significant differences for the different material combinations (**Table 1)**. Thus, image resolution is comparable throughout the eSee-Shell surface and most *in vivo* imaging experiments that can be performed using typical neuroimaging tools can be performed through eSee-Shells.

**Table 1:** Point spread function full-widths at half maximum.

|  | Lateral FWHM | Axial FWHM |
| --- | --- | --- |
| Glass coverslip | $0.99 \pm 0.09$ (n = 7) | $16.01 \pm 0.67$ (n = 7) |
| PET | $0.99 \pm 0.04$ (n = 7) | $16.10 \pm 0.47$ (n = 7) |
| PET + Parylene-C | $0.978 \pm 0.03$ (n = 20) | $15.54 \pm 0.44$ (n = 10) |
| PET + PEDOT:PSS + Parylene-C | $1.00 \pm 0.08$ (n = 22) | $16.05 \pm 0.94$ (n = 12) |

### eSee-Shell *in vivo* performance

The eSee-Shells were chronically implanted on eight mice (one C57BL/6, six Thy1-GCaMP6f^38^, and one Cux2-CreERT2^39^;Ai163^40^). **Figure 3a** shows an example of a Thy1-GCaMP6f mouse implanted with an eSee-Shell. The performance of the PEDOT:PSS ECoG electrodes *in vivo* was evaluated by tracking the magnitude and stability of the electrode impedances over time after implantation. As shown in **Figure 3b** for all ten electrodes in one eSee-Shell, the average impedance (after the removal of a single outlier) one week after implantation was 81.4 ± 53.4 kΩ at 1 kHz, which remained relatively stable for more than 100 days. We observed that low channel impedances (< 1 MΩ) generally correlated with good signal quality, however, some channels with high impedances were found to have good signal quality and some low-impedance channels had poor signal quality, indicating channels could not be categorized as working or non-working based on channel impedance alone. Hence, an electrode was defined as non-working if either of the following attributes were observed: 1) the ECoG had low-frequency oscillations with power greater than 150% of that of a low impedance electrode signal; or 2) the recordings contained significant 60 Hz line noise. We also tracked the number of working electrodes in eight implanted eSee-Shells for up to 150 days after implantation. On average, the implanted eSee-Shells had seven working electrodes (**Fig. 3c**). One implant had nine working electrodes for 256 days post-implantation, the longest duration assessed. Thus, eSee-Shells are robust and stable interfaces for measuring surface field potentials.

**Figure 3:**
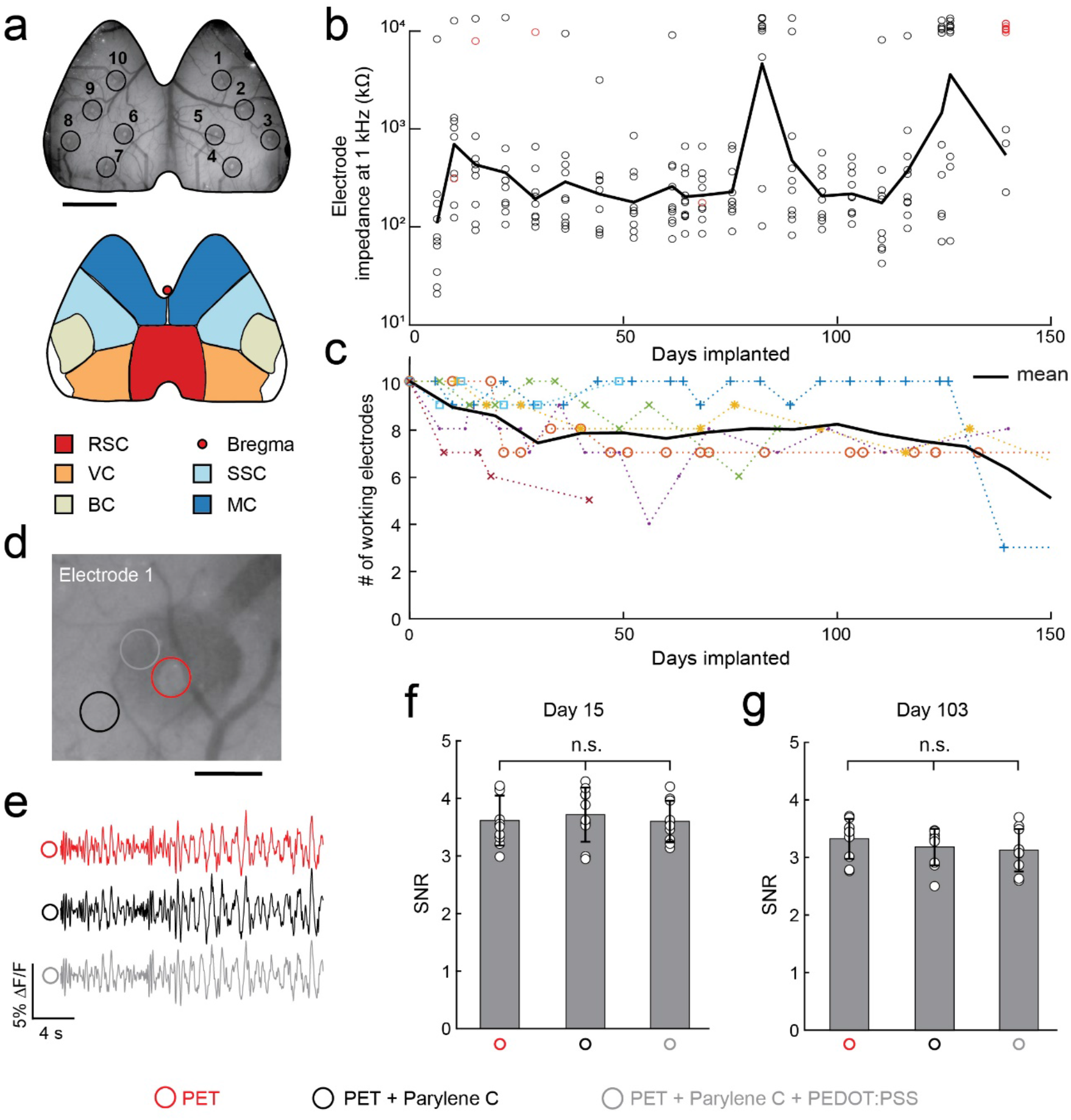
Chronic implantation of eSee-Shells: **(a)** Top: Bright-field image of a Thy1-GCaMP6f mouse brain implanted with an eSee-Shell, 7 weeks post-implantation. Circles and numbers indicate the position of each electrode. Scale bar indicates 2 mm. Bottom: Simplified version of the mouse brain atlas for the areas under the brain window. Specified areas are: Retrosplenial (RSC), Visual (VC), Barrel (BC), Somatosensory (SSC), and Motor (MC) Cortices. **(b)** Impedance of all 10 electrodes for an implanted eSee-Shell over time. Black and red circles indicate the impedance of working electrodes and non-working electrodes, respectively. **(c)** Number of working electrodes for different implanted eSee-Shells (n = 8) over time. Each color indicates one implant. Black line indicates the average number of working electrodes across all animals. **(d)** Fluorescence image of an electrode on an implanted eSee-Shell. Each circle indicates a different region of interest (ROI) of the transparent implant with different layers of material analyzed in panels **e-g**. Scale bar indicates 250 μm. **(e)** Representative ΔF/F traces for each of the ROIs indicated in panel **d**. **(f-g)** Quantification of signal-to-noise ratio (SNR) for similar ROIs indicated in **d** for all electrodes at 15 and 103 days after the implantation.

The eSee-Shell window over the brain has different layers of material at different locations. Therefore, we tested whether the multiple layers affected Ca^2+^ imaging. Much of the window area is the PET substrate covered by insulating Parylene-C, as shown by the black circle in **Figure 3d**. The regions along the interconnects have both PEDOT:PSS and Parylene-C on the PET substrate (**Fig. 3d**, grey circle). At the electrode contact sites, the Parylene-C and PEDOT:PSS are etched to leave the bare PET substrate (**Fig. 3d**, red circle). The Ca^2+^ signals obtained through these different material stacks were qualitatively similar, with the same activity patterns, as expected when imaging nearby ROIs (**Fig. 3d, e**). We calculated the signal-to-noise ratio (SNR) in sets of three ROIs under or in the vicinity of each of the ten electrodes on day 15 and 103 after implantation. On the same day, there was no significant difference (one-way ANOVA, F(2,9) = 0.22, *p* = 0.800 and F(2,9) = 0.87, *p* = 0.429 for day 15 and 103, respectively) in the SNR between the ROIs corresponding to the different material stacks (**Fig. 3f,g**).

### Measuring cortex-wide oscillatory activity under Ketamine anesthesia

Ketamine/Xylazine anesthesia generates brain-wide low-frequency oscillations in the mouse^41^. In Thy1-GCaMP6f mice (**Fig. 4a**), both the ECoG and Ca^2+^ signals reveal the expected low-frequency oscillations across the cerebral cortex under Ketamine anesthesia (**Fig. 4a,b**). Wide-field Ca^2+^ imaging reveals spatial dynamics, with strong periodic traveling oscillations originating in the anterior cortex and terminating in the posterior regions, which were coincident with large amplitude low frequency oscillations observed in the ECoG (**Supplementary Videos 1 and 2**, **Fig. 4c**). The dominant frequency power for both modalities was between 1 and 2 Hz with the peak power at ~1.5 Hz (**Fig. 4d,e**). At low frequencies, the two modalities are highly correlated (**Fig. 4g**). Under anesthesia, the ECoG has a second peak between 2.5 and 4 Hz that is not present in the Ca^2+^ signal. In addition to the low-frequency band, the Ca^2+^ signal exhibits increased power at 5-6 Hz, which is attributed to heartbeat artifacts^1^. Most importantly, ECoG samples higher frequency activity that cannot be captured by Ca^2+^ imaging (**Fig. 4f**). Although ECoG electrodes capture signals with higher frequencies, Ca^2+^ imaging provides more spatially accurate information about brain activity. Thus, the two modalities in eSee-Shells provide complementary datasets.

**Figure 4:**
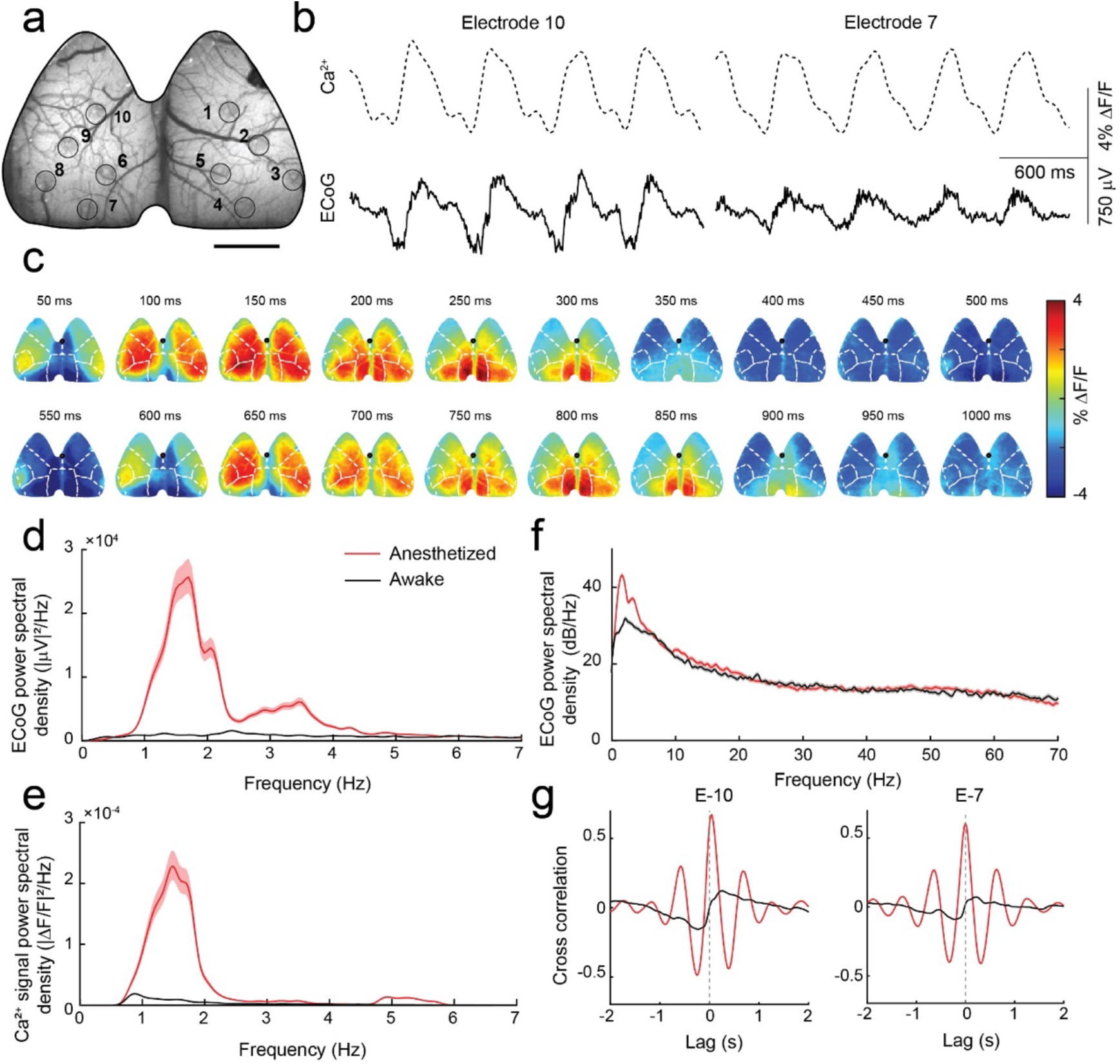
Comparison of ECoG and Ca^2+^ signals in an anesthetized and awake mouse: **(a)** Bright-field image of a Thy1-GCaMP6f mouse brain implanted with an eSee-Shell. Circles show the cortical locations of the ECoG electrodes, identified with numbers. Scale bar indicates 2 mm. **(b)** Simultaneously acquired ECoG from selected electrodes and Ca^2+^ signals from the area under the electrode during Ketamine-induced oscillations. Signals are recorded from the same implanted brain as in **a**. **(c)** Wide-field oscillations of Ca^2+^ signals over a 1 s period under Ketamine anesthesia. **(d,e)** Power spectral density (PSD) of the ECoG and Ca^2+^ signal from electrode 10 in awake and anesthetized states in the low frequency (1-7 Hz) range. Shaded areas represent 95% confidence intervals. **(f)** PSD of the ECoG from electrode 10 in the range of 1-70 Hz during both the anesthetized and awake state. **(g)** Cross-correlation between simultaneously recorded ECoG and Ca^2+^ signal under the same electrode in awake and anesthetized states for two different electrodes. The awake/anesthetized legend in **d** applies to panels **d-g**.

### Multimodal recording of sensory stimulus-evoked responses across the cortex

We measured mesoscale cortical dynamics in response to sensory stimuli using simultaneous ECoG and Ca^2+^ imaging. We refer to the ECoG signal from electrode *x* as “E-*x*”, and the Ca^2+^ signal beneath that electrode as “C-*x*”. In response to stimuli delivered to the whiskers, highly localized and spatially correlated ECoG and Ca^2+^ signals were observed (**Supplementary Video 3**). The peak Ca^2+^ signal response to left whisker stimulation was localized to the right barrel cortex (**Fig. 5a**), with a smaller response in the right motor cortex, consistent with previous studies of whisker-evoked spatial dynamics^42–44^. The Ca^2+^ signal beneath the electrode located in the contralateral barrel cortex (C-3) increased at ~50 ms post-stimulus and peaked around 150 ms, while no activity was detected in the Ca^2+^ signal beneath the corresponding ipsilateral electrode (C-8). Similarly, the ECoG electrode nearest to the right barrel cortex (E-3) captured the strongest average response to whisker deflection (−56 μV at 40 ms post-stimulus). The more anterior electrodes (E-1 and E-2) along the right sensorimotor cortex had detectable N1 and P2 peaks. Across mice, the same response pattern was observed for the N1 and P2 peaks at E-1, E-2, and E-3, highlighting the consistency of the evoked potentials across implantations (**Fig. 5c-e**).

**Figure 5:**
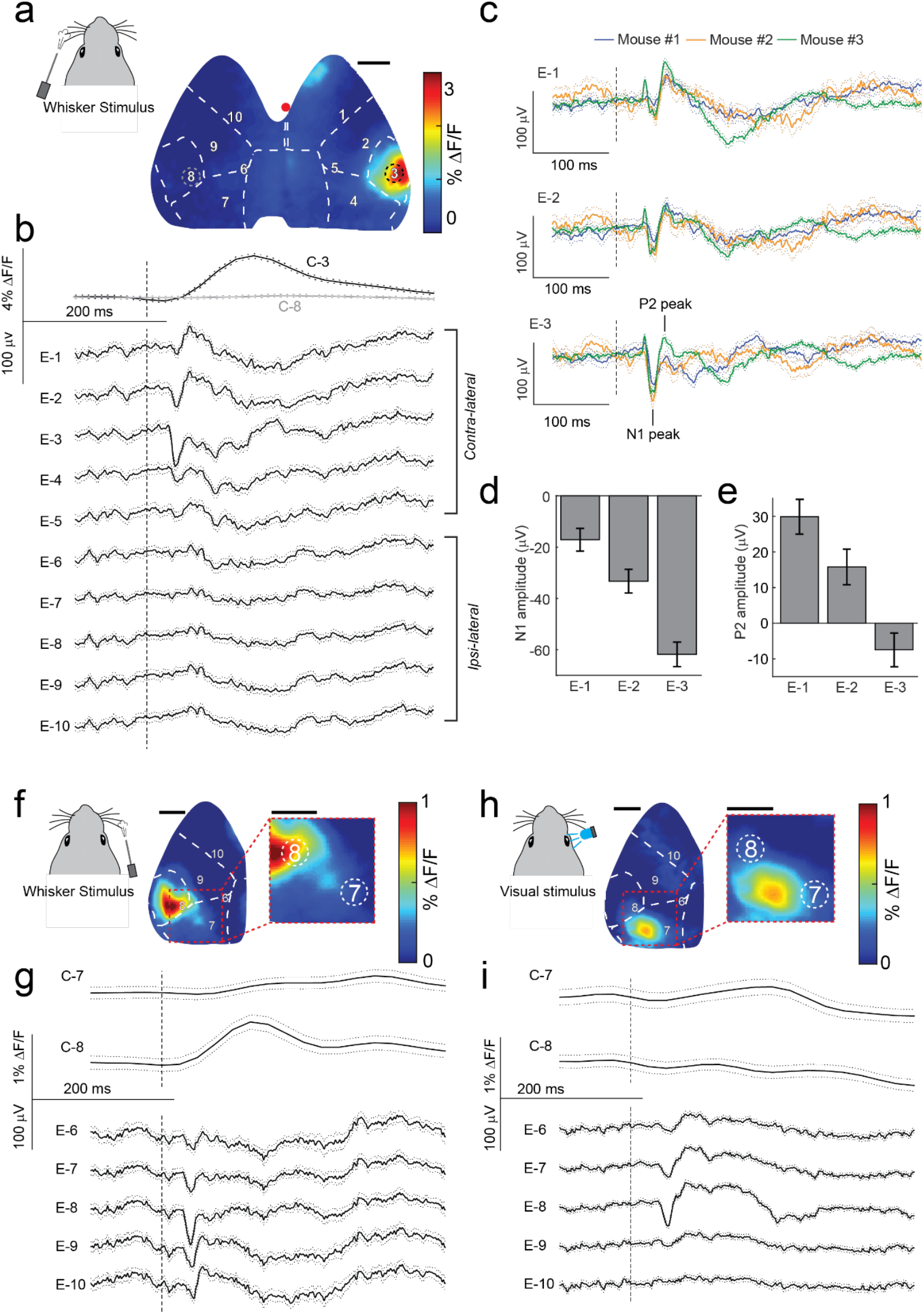
Measuring sensory evoked responses: **(a)** Spatial plot of the peak Ca^2+^ signal in response to whisker stimulation (n = 141 trials). **(b)** Ca^2+^ signals from ROIs indicated in **a** and ECoG across the cortex. **(c)** ECoG response of contra-lateral sensorimotor electrodes to whisker stimulation from three mice. **(d,e)** N1 and P2 amplitudes across all trials and mice (n = 272 trials) from contra-lateral sensorimotor electrodes in response to whisker stimulus. **(f)** Spatial plot of peak Ca^2+^ signal in response to whisker stimuli (n = 68 trials). **(g)** Ca^2+^ signal in ROIs indicated in **f** and ECoG of contra-lateral electrodes in response to stimuli. **(h)** Spatial plot of peak Ca^2+^ signal in response to visual stimuli (n = 174 trials) in the same mouse as **f**. **(i)** Ca^2+^ signal in ROIs indicated in **h** and ECoG on contra-lateral electrodes in response to stimuli. Vertical dashed lines indicate stimulus onset times. All confidence intervals were calculated using the jackknife standard deviation method. All scale bars indicate 1 mm.

We next compared the responses to whisker and visual stimulation in the same mouse. We observed a correspondingly similar response to right whisker stimulation as above for the left whisker stimulation. The peak ECoG and Ca^2+^ responses were localized to E-8 and C-8, respectively, corresponding to the left barrel cortex (**Fig. 5f,g**). However, visual stimulation evoked a peak ECoG response between E-7 and E-8 and a Ca^2+^ response between C-7 and C-8 (**Fig. 5h,i**). In response to whisker stimulation, the ECoG in the contralateral sensorimotor areas (E-8, E-9, and E-10) had prominent N1 and P2 peaks, while E-9 and E-10 were unresponsive to the visual stimulus. However, the latency and amplitude of the N1 peak differed between the two stimuli. Additionally, the P2 peak in response to visual stimuli was more prominent than for whisker stimulation. The sharp N1 and broad P2/N2/P3 complex are common features of mouse visual evoked potentials^45–47^. These experiments highlight the capability of eSee-Shells to record localized multimodal signals corresponding to activation of discrete sensory systems, illustrating the potential of these devices for studying and elucidating mechanisms underlying cortical sensory processing.

### Single-cell imaging through the ECoG electrodes

As a complement to the wide-field Ca^2+^ signals, cellular-resolution Ca^2+^ imaging was performed through the electrodes in eSee-Shells implanted on transgenic mice sparsely expressing GCaMP6s in layers 2/3 of the cortex (**Fig. 6a-c**). The sparse expression of GCaMP6s enables single-cell imaging at any location within the eSee-Shell field of view, while simultaneously acquiring ECoG signals across the cortex. Bright-field images of the brain around two electrodes are shown in **Figure 6d**. Individual neurons were detected using computational algorithms^48^ in the vicinity of two electrodes (C-7 and C-10), as well as underneath the electrodes and interconnects (**Fig. 6e**). The Ca^2+^ signal quality of the neurons under the electrodes was similar to the neurons outside the electrodes. **Figure 6f** illustrates the Ca^2+^ signal from a small subset of neurons imaged in the area near the two electrodes.

**Figure 6:**
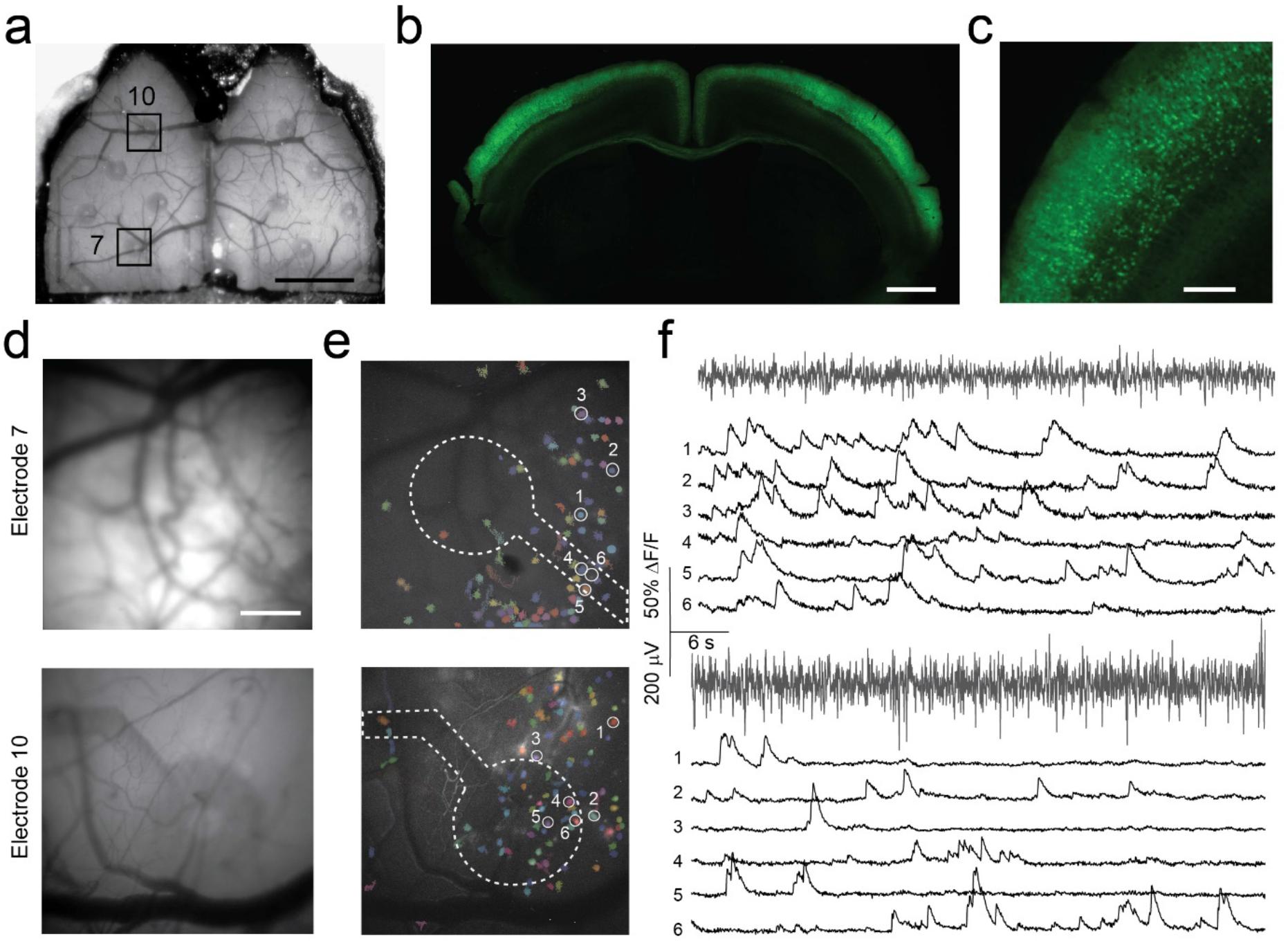
Single-cell imaging under and near transparent ECoG electrodes: **(a)** Wide-field image of a Cux2-CreERT2;Ai163 double transgenic mouse brain implanted with an eSee-Shell. ROIs around electrodes 7 and 10 are identified with boxes. Scale bar indicates 2 mm. **(b)** Coronal section of a Cux2-CreERT2;Ai163 double transgenic mouse brain expressing sparse GCaMP6s after being treated with tamoxifen. Scale bar indicates 1 mm. **(c)** Zoomed in region from **b** showing single neurons. Scale bar indicates 200 μm. **(d)** Mean image of a 250 s recording of Ca^2+^ fluorescence in ROI 7 (top) and 10 (bottom), respectively. The same order applies to panels **e** and **f**. Scale bar indicates 300 μm, with the same scale for panels in **e**. **(e)** Maximum intensity projection of the ΔF for the recordings in **d**, overlaid with the identified cells in the same ROI. Dashed lines show the electrode and interconnect location. Circles and numbers identify selected neurons. **(f)** ECoG signals and ΔF/F traces of selected neurons in the ROI.

### Multimodal, multiscale measurement of behavioral state-dependent cortical responses to sensory stimuli

Multiple studies have shown the effect of different behavior states on the brain’s response to a sensory stimulus^49–52^. Multimodal recordings using eSee-Shells not only enables observing these brain responses in high temporal and spatial resolution, but also allows study of state-dependent effects at different spatial scales simultaneously.

Here, we demonstrate these capabilities by segregating whisker stimulus trials into two categories based on whether the mouse was in an active whisking or quiescence state prior to the stimuli. At the mesoscale, the average Ca^2+^ signal response across the entire cortex shows distinct dynamics depending on whisking or quiescence. (**Supplementary Video 4**, **Fig. 7a-d**).

**Figure 7:**
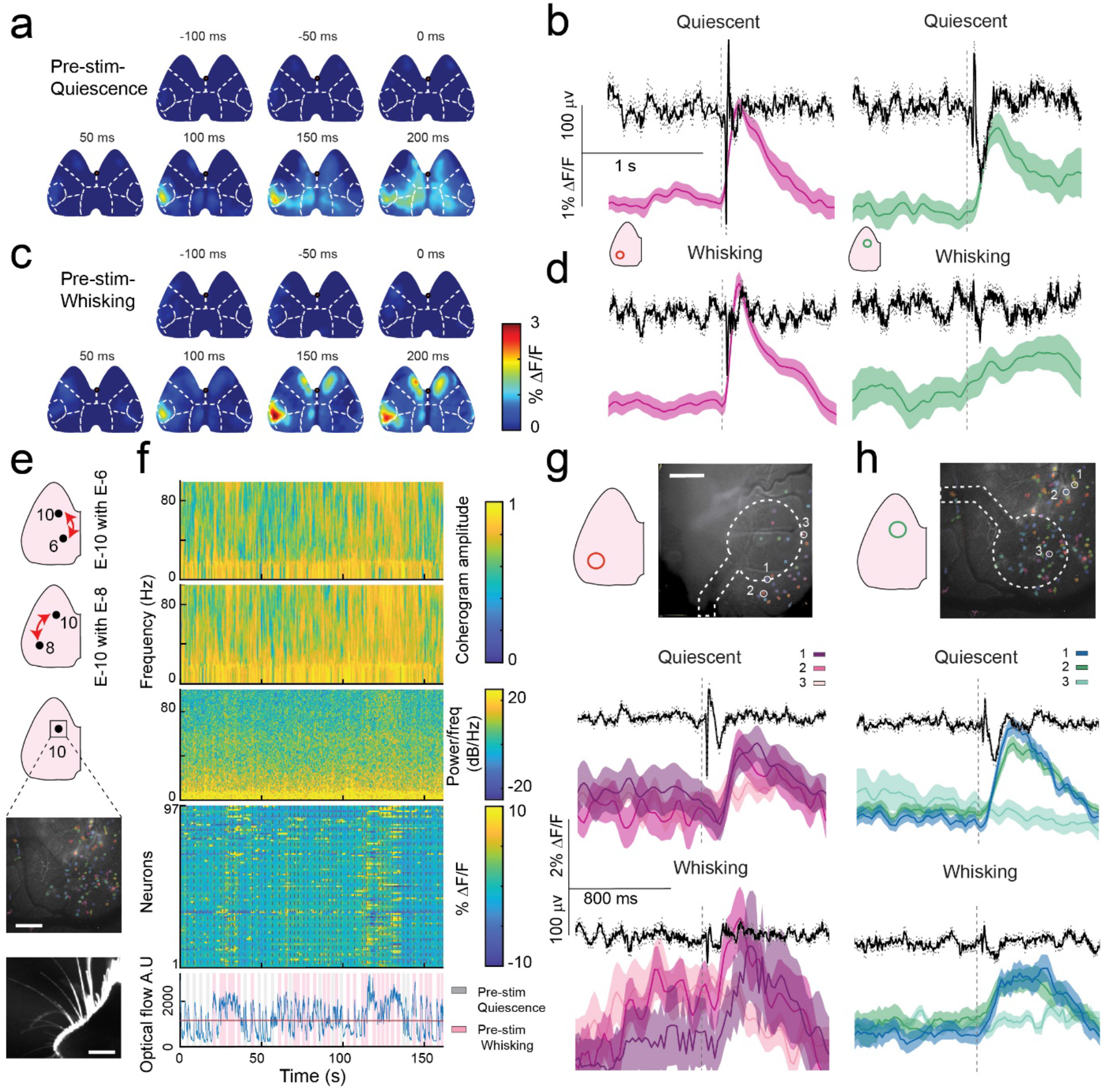
Multimodal, multiscale measurement of behavioral state-dependent cortical response to sensory stimuli: **(a)** Average mesoscale Ca^2+^ signals from a Thy1-GCaMP6f mouse brain throughout 300 ms in response to a 100 ms whisker puff during quiescent trials (n = 45). **(b)** Average ECoG (black) and Ca^2+^ signal color-coded from the specified ROIs in the quiescent trials. Averaged trials are the same as those used for panel **a**. **(c)** Average mesoscale Ca^2+^ signals in response to the whisker puff stimulus during whisking trials (n = 20). **(d)** Average ECoG and Ca^2+^ signal from the specified ROIs in the quiescent trials. Axis scales are the same for all subpanels in **b** and **d. (e)** From top to bottom: position of electrodes 10, 6, and 8. Middle: the ROI around electrode 10. Second to bottom: single neurons detected under electrode 10. Scale bar indicates 300 μm. Bottom: a snapshot of the behavioral camera video used to analyze the whisking state of the mouse. Scale bar indicates 4 mm. **(f)** Top: Coherograms of E-10 with E-6 and E-10 with E-8 from an eSee-Shell implanted on a Cux2-CreERT2;Ai163 mouse for a 150 s recording. Middle: Spectrogram of E-10 in the same period as the top. Second to bottom: ΔF/F signals from individual neurons shown on the left. Bottom: Optical flow of the whisker video as a measure of the whisking state. Horizontal line indicates the threshold for counting a state as whisking. Gray columns are quiescent trials (middle of the column), while pink columns indicate whisking trials. **(g, h)** Top-left: Position of the electrode (ROI). Scale bar indicates 300 μm. Top-right: ROI and the identified neurons. Middle and bottom: Average ECoG of the specified electrode and Ca^2+^ signals of selected neurons in that ROI segregated based on the whisking state before the whisker puff stimulus. Axis scales are the same for all plots in **g** and **h**.

For C-8 and C-10, robust peaks in the Ca^2+^ signals were observed in the barrel cortex area (C-8) in response to the stimulus in both behavioral states, with a significantly higher peak response of 2.1 ± 0.2% ΔF/F observed when whisking (n = 20) as compared to 1.4 ± 0.1% ΔF/F (n = 45) peak response when quiescent (*p* = 0.0026, Wilcoxon rank sum test, **Fig. 7b,d** left panels). In contrast, a sharp peak in the Ca^2+^ signal was present under the ECoG electrode in the motor cortex area (C-10) only when quiescent. Whisking produced higher baseline activity than when quiescent. When the mice were whisking the peak response in the Ca^2+^ signal was 0.7 ± 0.3% ΔF/F at 600 ms after the stimulus. However, when quiescent, the peak response of 0.7 ± 0.1% ΔF/F occurred 200 ms after the stimulus.

Qualitatively, the ECoG recordings from E-8 and E-10 also showed distinct responses based on the whisking state. When quiescent, prominent N1 and P2 peaks were observed in both the electrode above the barrel cortex regions (E-8) and the electrode above the motor cortex (E-10). Both peaks were diminished in the whisking state. In the electrode above the barrel area (E-8), the N1 response was −135 ± 18 μV at 40 ms after stimulus presentation when whisking, which was significantly smaller than −264 ± 13 μV when quiescent (*p*=1.2×10^−5^, Wilcoxon rank sum test). In the motor cortex (E-10), an N1 peak of −94 ± 14 μV was observed when the mouse was quiescent (**Fig. 7b,d**). No distinct N1 peak above baseline activity was observed when actively whisking. These state dependent responses to whisker stimulation agree with previous findings using voltage-sensitive dyes^42^.

We performed the same state-dependent segregation of whisker-stimulus responses in a Cux2-CreERT2;Ai163 mouse (**Fig. 7e-h**). Periods of sustained whisking activity resulted in synchronous activation of large populations of neurons. Correspondingly, an increase in power of the ECoG at higher frequencies was evident as well as higher coherence at the high frequencies with both proximal and distal electrodes. **Figure 7g** and **h** show average Ca^2+^ signals of 3 neurons near electrodes at the barrel and motor cortices, respectively, along with simultaneously measured ECoG. We observed similar state-dependent ECoG dynamics, consistent with our previous experiment in the Thy1-GCaMP6f mouse (**Fig. 7b-d**). However, compared to mesoscale Ca^2+^ signals, heterogeneous cellular responses occurred to sensory stimulus presentation during both behavioral states (**Fig. 7e,f**). Despite the higher magnitude ECoG responses during the quiescent trials, the mesoscale Ca^2+^ signals from the area underneath the electrodes had a lower magnitude compared to the whisking trials. This discrepancy is most likely due to the inherent difference between the ECoG and Ca^2+^ signals. The magnitude of the ECoG depends not only on the activity of individual neurons but also on other parameters, such as the neurons’ orientation and the synchrony of the activity^53^. Although the neural activity generally increased during whisking, this activity is less synchronized, causing the reduction of the magnitude of the recorded ECoG^54,55^.

## DISCUSSION

Previous studies demonstrated the advantage of transparent electrodes by performing simultaneous 2-photon imaging and ECoG acquisition^20–22,25,26,28^. However, the implants covered small regions of the mouse brain and hence, the recorded ECoG only captured neuronal activity in the vicinity of the imaged neurons. Here we report simultaneous neuronal Ca^2+^ imaging and ECoG recordings over much of the mouse dorsal cerebral cortex using eSee-Shells. Our eSee-Shells allow either mesoscale imaging across the entire FOV or random-access cellular level imaging of small FOVs throughout ~45 mm^2^ of the cerebral cortex. These advances in multimodal, multiscale sensing were made possible by the development of transparent, flexible and low impedance PEDOT:PSS inkjet-printed ECoG arrays. Further, these large-scale, dual recordings can be maintained over 100 days. To our knowledge, this is the first demonstration of truly long-term, flexible neural devices than can be implanted over much of the dorsal cortex of the mouse.

ECoG is a highly valuable and widely used electrophysiology modality that allows for minimally invasive monitoring of population activity across large regions of the cortex. Yet, the underlying contributions to ECoG from different cell types are not fully understood. Similarly, mesoscale Ca^2+^ imaging using transgenic animals broadly expressing Ca^2+^ indicators have revealed new insights into how large-scale cortical activity is modulated in a variety of behaviors, though the contribution of different sources to mesoscale Ca^2+^ signals are yet to be determined^4,5,8,42,56^. This study highlights the effect of both behavioral and brain state on ECoG dynamics. Combining ECoG with high resolution cellular scale imaging as done here, can in the future reveal the cellular mechanisms of ECoG and mesoscale Ca^2+^ signals. While Ca^2+^ dynamics are limited to low frequencies, in the future, the eSee-Shells could be implanted with transgenic mice expressing genetically encoded voltage indicators^57,58^ to investigate the contribution of specific cell types to the ECoG. Similarly, widespread Ca^2+^ imaging is becoming increasingly popular. Expressing Ca^2+^ indicators in specific cells and cellular compartments^59–61^ would allow determination of distinct cell and sub-cellular contributions to mesoscale Ca^2+^ signals and ECoG.

There are some limitations to our current approach that can be improved upon in future versions of the eSee-Shells. We patterned a relatively small number of electrodes in the polymer skulls. A primary limitation on this channel count using our approach is the ‘neck’ region of the device located posterior to the cranial window. This neck is 4 mm at its narrowest. While inkjet printing is highly customizable, compared to traditional photolithography processes, the achievable interconnect pitch is much larger (on the order of ~100 μm), limiting the number of interconnects that can be routed through this region. In the future, we can explore a multilayered printing approach in the ‘neck’ region to build eSee-Shells with a larger number of electrodes. Furthermore, because inkjet printing relies on a digital pattern input rather than a physical pattern (such as a photolithography mask), the number and placement of the ECoG electrodes can be rapidly tailored to target specific brain regions in different experiments. The development of semiconducting inks^62^ and multilayered inkjet printing^63^ could enable inkjet printing of multiplexed circuits for high channel count transparent ECoG devices. The multimodal recordings capabilities of the eSee-Shells can in the future be extended to freely behaving animals, by engineering miniaturized electronic interfaces along with existing optical devices^33^.

## METHODS

### Device Fabrication

#### Substrate laser-patterning and cleaning

50 μm thick Polyethylene terephthalate (PET) substrates (Melinex 462, Dupont Teijin) were laser patterned using a free-standing laser cutter (PLS6.150D, Universal Laser Systems, Inc.) to create the electrode array outline and alignment features used for bonding the array into the window frame. The substrates were then sprayed with isopropyl alcohol (IPA), acetone, and distilled water (DW) to remove large contaminants resulting from the laser patterning, followed by sonication for 5 minutes each in methanol and a detergent solution (2% Micro-90 in DW). Finally, the detergent solution was sprayed off with DW, and substrates underwent a final 5-minute sonication in pure DW.

#### Inkjet printing of silver and PEDOT:PSS features

An inkjet printer (LP50 Pixdro, Meyer Burger Technology Ltd) was used for patterning silver (Silverjet DGP-40LT-15C, ANP Inc.) and PEDOT:PSS (Orgacon™ IJ-1005, Agfa-Gevaert N.V.) solutions. Briefly, for printing silver patterns, the platen temperature was set to 65°C and the print speed to 75 mm/s. Silver features were printed in 2 layers to improve conductivity. Silver patterns were then sintered on a hot plate for 2 hours at 100°C. Prior to patterning PEDOT:PSS features, the PET surface energy was modified by brief exposure to argon plasma (12 W, 400 mTorr, 100 sccm Ar, 25 s; STS 320 RIE, Surface Technology Systems). PEDOT:PSS features were then inkjet printed with the platen temperature set to 65°C with a print speed of 100 mm/s. PEDOT:PSS films were subsequently cured on a hot plate for 15 minutes at 130°C.

#### Parylene deposition and patterning

Printed bond pads, and the back surface of the PET, were masked by polyimide tape (Kapton®, Dupont™) to prevent Parylene deposition. Electrode arrays were then coated in ~2.5 μm of Parylene-C (Labcoater 2, Specialty Coating Systems Inc.). Polyimide tape was adhered temporarily to glass microscope slides and contact opening shadow masks were laser-patterned (PLS6.150D, Universal Laser Systems, Inc.) in the tape. Shadow masks were then manually aligned and adhered to the electrode arrays. Contact site patterns were etched into the arrays through reactive-ion etching (Advanced Vacuum Vision 320 RIE, Plasma-therm).

#### Final eSee-Shell assembly

Electrically conductive adhesive transfer tape (ECATT, 9703, 3M) was used to bond the inkjet-printed bond pads and the bond pads on custom interface PCBs (JLCPCB). The seam between the PET substrate and the PCB was then sealed with cyanoacrylate adhesive (3M liquid superglue). Electrode arrays were then bonded to 3D-printed window frames as described previously^11,33^.

### Benchtop testing

#### Electrochemical impedance spectroscopy

Electrochemical impedance spectroscopy was performed with a potentiostat (Gamry Interface 1010, Gamry Instruments Inc.) in room temperature phosphate buffered saline (PBS, # D1283, Sigma Aldrich). A platinum wire was used as a counter electrode. Once the electrodes were submerged in PBS, an open-circuit potential stability of <0.1 mV/S was reached after an equilibration period lasting ~5 minutes. A 50 mV root mean square AC voltage was used for impedance measurements, and measurement frequency was swept from 10 kHz – 0.1 Hz, with four measurements acquired per decade.

#### Point spread function measurement

Agar powder (Carolina Biological Supply Company) and sodium chloride (Fisher Chemical™) were dissolved (2.0:0.5:97.5% w/w) in boiling deionized water. While still heated, 1 mL of the agar solution was repeatedly aspirated with 200 μL of a 1:500 dilution of 0.1 μm yellow-green fluorescent nanobeads (FluoSpheres®, Invitrogen) to evenly distribute beads in the agar solution. The solution was then pipetted onto a standard glass microscope slide (Plain Glass Micro Slides, Fisherbrand™) and the appropriate substrate, either a glass coverslip (Square #1½ Cover Glass, Corning) or an electrode array, was pressed on top to complete the sample. The edges were sealed with silicone elastomer (Kwik-Sil, WPI Inc.) and stored at 4°C to prevent dehydration. A confocal microscope (C2, Nikon) with a 40X, 0.6 numerical aperture (NA) air objective was used to image bead point spread functions (PSFs). A 488 nm laser was used to excite nanobead fluorescence, and identical laser power and detector settings were used for all samples. A 30 μm pinhole size was used, and z-sections were imaged at focal planes from 20 μm below to 20 μm above each bead in 0.5 μm intervals. Each final image was produced by averaging 8 separate images. The lateral and axial intensity profiles for each bead PSF were extracted in FIJI^64^ followed by the use of custom scripts for fitting gaussian curves to each profile to extract FWHMs (Matlab, The Mathworks Inc.).

### Surgical Implantation

All animal experiments were approved and conducted in accordance with the University of Minnesota’s Institutional Animal Care and Use Committee (IACUC). eSee-Shells were implanted on a C57BL/6 mouse (JAX000664, Jackson Laboratories) and 6 Thy1-GCaMP6f transgenic mice (JAX024339, Jackson Laboratories) that express GCaMP6f in excitatory neurons in layers 2/3 and 5 of the cerebral cortex^38^. For single-cell imaging, an eSee-Shell was implanted on a double transgenic mouse Cux2-CreERT2;Ai163-GCaMP6s (032779, Mutant Mouse Resource and Research Center; Ai163, Allen Institute) which selectively expresses GCaMP6s in layer 2/3 excitatory neurons via administration of 75 mg/kg Tamoxifen for 5 days^39,40^.

Mice were anesthetized using a gas mixture of 95% oxygen and isoflurane (5% for induction, 1-3% for maintenance). Prior to surgery, mice were administered 2 mg/kg slow-release Buprenorphine (Buprenorphine SR-LAB, Zoopharm Inc.) and 2mg/kg Meloxicam (Loxicom®). The depth of anesthesia was monitored by toe pinch response every 15 minutes throughout the surgical procedure. Eyes were covered with sterile eye ointment (Puralube, Dechra Veterinary Products) and the scalp disinfected with alternating betadine and 70% ethanol washes. The surgical procedure began with excision of the scalp, followed by removal of the fascia. A self-tapping bone screw (F000CE094, Morris Precision Screws and Parts) was implanted on the occipital bone, 2-3 mm inferior to the interparietal bone and mediolateral from to lambda as both an anchor and as reference electrode for the ECoG electrodes. A stainless-steel conductive thread (#640, Adafruit Industries) was tied to the screw to facilitate electrical access to the screw during ECoG acquisition. A craniotomy was performed to remove a flap of skull that matched the geometry of the implant window. Care was taken to ensure the dura remained intact. Gauze soaked in sterile saline was used to clear any bleeding from the bone removal procedure. The eSee-Shell was mounted on a custom holder attached to a stereotaxic arm, and carefully lowered into position over the craniotomy. A small amount of tissue adhesive (Vetbond™, 3M) was applied to the sides of the implant to fix it to the skull followed by application of dental cement (Metabond, Parkell Inc.) around the implant and skull screw. The titanium head plate was fastened to the eSee-Shell frame using flat head screws (3/32 inch, 0-80). The PCB tail of the electrode array was fixed to the head plate using tissue adhesive. In the end, the whole assembly was further secured by covering the connection sites with dental cement. A 3D-printed protective cap was then attached to the head plate. The mice recovered to an ambulatory state on a heating pad and then were returned to a clean home cage.

### Head-fixed imaging and electrophysiological recordings

#### Head-fixation

All imaging and electrophysiology experiments were conducted on a custom-built treadmill setup wherein head-fixed mice were able to locomote on a rotating disk^65^. Mice were acclimatized to handling and the treadmill by being allowed to explore the treadmill without restraint prior to experiments.

#### ECoG Acquisition

A 128-channel amplifier head stage (RHD 128-Channel Recording Headstage, Intan Technologies, LLC.) was connected to the eSee-Shell after head fixation through the custom adapter PCB. The head stage was connected to an interface board (RHD USB interface board, Intan Technologies, LLC.). ECoGs were high-pass filtered with a 0.7 Hz cut-off frequency and recorded at 20 kHz.

#### Ca^2+^ imaging

The Ca^2+^ signals were imaged using an epi-fluorescence stereo-zoom microscope (MZ10, Leica) equipped with a high-speed imaging camera (Orca 4, Hamamatsu Inc.). Images were acquired at 40 Hz (512×512 pixels, 8-bit output depth) at 1.6X magnification. Cellular resolution imaging was performed at 12X magnification using the same setup.

#### Data synchronization

Ca^2+^ imaging and electrophysiology data streams were synchronized by sending a transistor-transistor logic (TTL) pulse at the acquisition of each image frame to the Intan USB interface. Exposure time, ECoGs, and stimulus presentation trigger signal were recorded at 20 kHz.

#### Ketamine/Xylazine anesthesia

In a Thy1-GCaMP6f mouse implanted with an eSee-Shell, ECoG and Ca^2+^ signals were recorded first in the spontaneous awake state. In this experiment the mouse was initially anesthetized using 1% mixture of isoflurane and oxygen, and after 2 minutes injected with a cocktail of Ketamine/Xylazine (100/10 mg/kg, intramuscular), followed by the dual recordings.

#### Sensory stimulus presentation

Visual and whisker stimuli were presented to awake mice head-fixed on the custom treadmill. For whisker stimuli, a compressed air supply was connected to a 24 gauge blunt stainless steel needle through a solenoid valve. The needle was placed so that the air puff stimulated the whiskers in the anterior-posterior direction and was not directed at the whisker pads. The solenoid valve was controlled using a microcontroller (Arduino Uno, Arduino LLC) to deliver 100 ms air puffs at randomized intervals between 4-6 s. Each experiment lasted 5 minutes and consisted of ~50 stimuli. Timing of sensory stimuli was logged by sending a TTL pulse from the microcontroller to a digital input channel in the Intan data acquisition board.

Visual stimuli were presented using a 7 inch computer monitor screen placed ~2 cm in front of the mouse, perpendicular to the visual axis of the right eye. A shroud ensured light from the monitor did not enter the wide-field microscope. Ten 50 ms white flashes were presented at 1 Hz in each trial^66^, and trials were separated by a random length of time lasting 4-8 s. A photoresistor taped to the corner of the monitor recorded the visual stimuli signal.

#### Whisking behavior recording

Behavioral imaging was conducted using a high-speed monochrome camera (Blackfly® S USB3, FLIR). Whisker movement was recorded at 150 Hz. Two infrared LED lamps (48 LED IR Illuminator, OLSUS) illuminated the whiskers.

### Data analysis

All data analysis was carried out using in built functions and custom scripts written in MATLAB (The Mathworks Inc.), and specific MATLAB functions are placed in quotation marks.

#### Mesoscale imaging data analysis

The Ca^2+^ videos were motion corrected using the “dftregistration” function^67^. Then, the data was band-pass filtered in the time domain using a 4^th^ order FIR filter with 0.1/7 Hz cutoff frequencies. The ΔF/F was calculated by dividing the Ca^2+^ signal intensity of each pixel to the average of the intensity of that pixel over the recording time. To ease the large file size of the Ca^2+^ signals and to further reduce noise in the Ca^2+^ signal, we performed fast singular value decomposition (f-SVD) on the data using “compute_svd”^68,69^.

The average Ca^2+^ signals of the mouse brain in response to the sensory stimulus were calculated by recomposing the SVD matrices of the ΔF/F videos and binning them into 25 ms peri-stimulus bins for each trial. The frames within each 25 ms bin were averaged to approximate the average Ca^2+^ signal for that time period. Power spectra of mesoscale Ca^2+^ signals were calculated using a built in “pwelch” function with a Hanning window of 7.1 s with a 6.4 s overlap between windows. Maximum ΔF/F was defined as the maximum value of ΔF/F in a spontaneous recording. SNR was calculated as the ratio between the maximum ΔF/F to the variance of ΔF/F in the same recording.

#### Single-cell imaging data analysis

Suite2p^48^ was used to isolate and extract single-cell Ca^2+^ transients. Maximum intensity projections of single-cell Ca^2+^ videos were generated using Fiji and the MOCO plugin^70^. The extracted traces were corrected by removing the neuropil signal around each detected neuron. The resultant signal was filtered with a 4^th^ order FIR bandpass filter with lower cutoff frequency of 0.005 Hz and higher cutoff frequency of 8 Hz, then normalized by the average Ca^2+^ signal magnitude for each cell. Finally, up to the quadratic trend was removed from the signal using the built-in “detrend” function. The lower cutoff frequency was set close to zero to avoid any high-pass filter artifact^45^ on the low-frequency Ca^2+^ signal while removing the DC offset.

#### Electrophysiology data analysis

ECoGs were low-pass filtered with a 50^th^ order elliptic filter (cutoff frequency of 300 Hz), followed by down sampling by 9. Analog and digital inputs from stimulus presentation electronics were also down sampled and thresholded to find timepoints where stimuli occurred, referred to hereafter as trials. Power spectra and spectrograms of ECoGs from Ketamine/Xylazine anesthesia were calculated using “pwelch” function with a variable Hanning window depending on the range of frequencies analyzed. For recordings of sensory stimulus evoked potentials, trials in which the ECoG magnitude exceeded ± 400 μV within 1 s of the stimulus presentation were considered artifacts and were removed from further analysis. The mean and standard deviation of the ECoG from the remaining trials were then calculated using the jackknife standard deviation method^71^. The amplitudes of N1 and P2 peaks of whisker-puff event-related potentials (ERPs) were determined from the ECoG amplitude at 40 and 55 ms, post-stimulus presentation, respectively. Coherency was calculated by first segmenting the ECoGs into 1 s windows with 0.75 s overlap followed by computing the coherence using the multi-taper method^71^.

#### Whisking behavior analysis

To find instances of whisking, we measured the motion of the mouse’s whiskers and snout (**Fig. 7e**, bottom). The motion in the field of view was quantified using the “optical flow” function. A minimum threshold was manually set on the measured optical flow for each frame to detect moving states. The recorded whisker puff trials were then segregated based on the number of moving frames 100 ms (15 total frames) before the whisker puff onset. If the number was greater than 4, that trial was marked as an active trial. Otherwise, the trial was marked as a quiescent trial. The number of moving frames required for a trial to be labelled as active was chosen to cover at least half the time of one 20 Hz whisking cycle^72^. Responses during active and quiescent trials were then averaged separately.

### Histology

#### Tissue Fixation and Slicing

After experiments, the Cux2-creERT2;Ai163 mouse was fully anesthetized in 5% isoflurane and transcardially perfused with phosphate-buffered saline (P5493, Sigma Aldrich) followed by 4% paraformaldehyde (P6148, Sigma Aldrich). The brain was extracted and post-fixed in 4% PFA for 24 hours, and transferred to 30% sucrose (S0389, Sigma Aldrich) for 48 hours. The tissue was then flash frozen in 2-Methylbutane (O3551-4, Fisher Chemical™) on dry ice and kept at −80°C until sectioned. Using a cryostat, 40 μm coronal sections were thaw mounted on slides (71869, Electron Microscopy Sciences) and mounted with mounting medium (Vectashield H-1500, Vector Labs).

#### Imaging

Sections were imaged using an epifluorescent microscope (BZ-X710, Keyence). 4x images were acquired and stitched together into a mosaic using corresponding Keyence software.

## AUTHOR CONTRIBUTIONS

These authors contributed equally: Preston D. Donaldson, Zahra S. Navabi. P.D.D., Z.S.N., S.B.K., S.L.S., R.E.C., T.J.E., and L.G. contributed to the eSee-Shell design. Z.S.N., P.D.D., S.B.K., S.L.S., R.E.C., and T.J.E. designed experiments and wrote the manuscript. P.D.D. optimized the eSee-Shell fabrication and conducted benchtop characterizations. Z.S.N. and R.E.C. performed the animal surgeries. Z.S.N., P.D.D., and R.E.C. optimized surgical procedures, conducted the multimodal experiments, and performed analysis of *in vivo* data. S.M.L.F. performed histological analysis.

## ACKNOWLEDGEMENTS

SBK SLS and TJE acknowledge NINDS Award #R0NS111028. SBK acknowledges Brain Initiative Award R42NS110165. Microfabrication and PSF characterizations were performed at the Minnesota Nano Center, funded by NSF NNCI Award ECCS-2025124. PDD was supported by NSF IGERT Award DGE-1069104. SBK and TJE acknowledge P30DA048742.

## INSTITUTIONAL APPROVAL

All animal experiments described in this paper were approved by the University of Minnesota’s Institutional Animal Care and Use Committee (IACUC).

## DATA AVAILABILITY

Full numerical data shown in the figures are available as a supplementary dataset accompanying the article. Raw imaging datasets will be made available on request.

## LIST OF SUPPLEMENTARY MATERIAL

**Supplementary Video 1:** Awake vs anesthetized ECoG and Ca^2+^ signals.

**Supplementary Video 2:** ECoG band powers and Mesoscale Ca^2+^ dynamics in awake and anesthetized states.

**Supplementary Video 3:** Average ECoG and Ca^2+^ whisker stimulus evoked response.

**Supplementary Video 4:** Multimodal recordings during active whisking and quiescence.

